# Genes for tRNA recycling are upregulated in response to infection with Theiler’s mouse encephalitis virus

**DOI:** 10.1101/2021.08.05.455008

**Authors:** Mineaki Seki, Akihiko Komuro, Tatsuya Ishikawa, Masayuki Takahashi, Masayuki Nashimoto

## Abstract

The concept of tRNA recycling has recently emerged from the studies of ribosome-associated quality control. Therein tRNase ZS removes the 2′, 3′>p from the ANKZF1-cleaved tRNA and the subsequent TRNT1 action re-generates the intact tRNA. To know the roles of the tRNA recycling in vivo, we investigated how viral infection affects the tRNA recycling system by analyzing the mRNA levels of tRNase ZS and TRNT1. We found that both genes in HeLa cells are upregulated in response to infection of Theiler’s mouse encephalitis virus but not to that of an influenza A virus. Upregulation was also observed in cells infected with encephalomyocarditis virus with reduced efficiency. The levels of the IFN-β mRNA appeared to positively correlate with those of the tRNase ZS and TRNT1 mRNAs. The tRNase ZS gene may be regulated post-transcriptionally in the cells infected with Theiler’s mouse encephalitis virus.

## Introduction

tRNA recycling has recently found in the studies of ribosome-associated <u>q</u>uality <u>c</u>ontrol (RQC). The physiological roles are unclear and biochemical studies revealed that two enzymes, namely tRNase ZS (ELAC1) and TRNT1 are involved in this reaction ^1,2^. tRNase ZS is a paralog of tRNA maturation enzymes, tRNase ZL (or ELAC2), which cleaves pre-tRNAs at the junction of a discriminator nucleotide and a 3′ trailer sequence, enabling the subsequent 3′-terminal CCA addition ^3–5^. On the other hand, tRNase ZS has very limited pre-tRNA-cleaving activity compared with tRNase ZL ^6,7^. This property together with the fact that the tRNase ZS gene is dispensable for viability of at least one human cell line made us curious about its physiological role ^8^. Another tRNA maturation enzyme, TRNT1 (or the CCA-adding enzyme), catalyzes the addition of CCA nucleotides^9^. This activity is used not only for tRNA maturation but for tRNA surveillance and quality control ^10,11^.

RQC prevents cells from accumulating faulty proteins that can have cytotoxic properties by degrading the aberrant polypeptides ^12,13^. This process starts with ubiquitination of an aberrant nascent chain and ends with its proteasomal degradation after extracting it from a 60S ribosome-nascent chain complex (60S-RQC). This extraction relies on the peptidyl-tRNA hydrolase activity of ANKZF1, which can cleave the peptidyl-tRNA in the 60S-RQC at the junction of the 3′-terminal CCA nucleotides and the discriminator nucleotide to release the tRNA with a 2′, 3′-cyclic phosphate (2′, 3′>p) ^14–16^. The CCA-less tRNA with the 2′, 3′>p seemed to be resistant not only to exonuclease digestion but also to repair by TRNT1. This apparent dead-end situation and the enigma of role of tRNase ZS have been solved simultaneously by the discovery that tRNase ZS can remove the 2′, 3′>p from the ANKZF1-cleaved tRNA and that the subsequent TRNT1 action re-generates the intact tRNA ^1,2^.

Here, we investigated how viral infection affects the tRNA recycling system by analyzing the expression levels of tRNase ZS and TRNT1. We uncovered that that both genes are upregulated in response to infection of Theiler’s mouse encephalitis virus (TMEV) or the encephalomyocarditis virus (EMCV) but not to that of an influenza A virus (IAV).

## Materials and methods

### Cell culture

HeLa cells were cultured in complete DMEM media (DMEM media containing 10% fetal bovine serum and 100 U/mL penicillin-streptomycin) at 37 °C in a 5% CO_2_ incubator.

### Viral infection

HeLa cells were seeded and cultured in 24-well plates. After washing with phosphate-buffered saline, the cells were incubated in reduced (2% fetal calf serum) complete media containing virus for 2 h. Subsequently, the media was replaced by normal (10% fetal calf serum) complete media ^17^. The cells were then collected at the indicated time points and total RNA was extracted.

### Quantitative reverse transcription PCR (qRT-PCR) analysis

Total RNA was extracted using the RNAiso Plus (Takara Bio, Shiga, Japan). cDNA synthesis was performed using the ReverTra Ace qPCR RT Master Mix with gDNA Remover (Toyobo, Osaka, Japan) in accordance with the manufacturer’s protocol. mRNA levels were quantitated using the THUNDERBIRD SYBR qPCR Mix (Toyobo) and normalized against glyceraldehyde-3-phosphate dehydrogenase (GAPDH) mRNA levels. The fold-change in mRNA levels was calculated using the standard curve method. The primer pairs used are listed in Table 1.

**Table 1.**
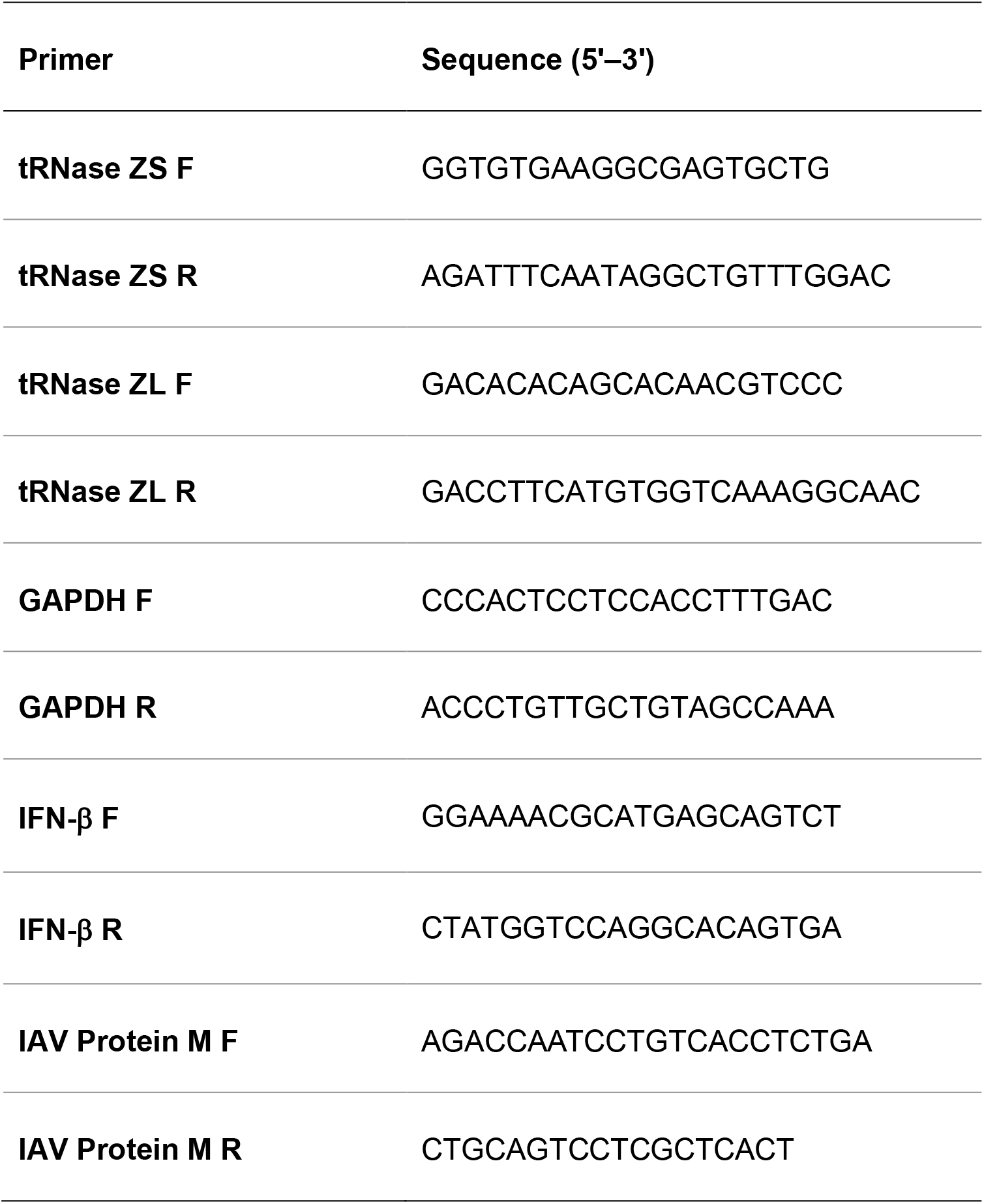
Sequences of primers used in this study.

### RT-PCR assays

Total RNA was extracted from IAV-infected cells using the RNAiso Plus kit. cDNA synthesis was performed using the ReverTra Ace qPCR RT Master Mix with gDNA Remover in accordance with the manufacturer’s protocol. RT-PCR was performed using the AmpliTaq Gold 360 Master Mix (Thermo Fisher Scientific, Waltham, MA, USA). The reaction mixture (20 μL) included the IAV protein M primer pair (4 μM) and cDNA template (6 ng) ^18^. PCR was performed for 36 cycles, and the products were separated on a 2% agarose gel.

### Immunoblotting

HeLa cells were suspended in NETN buffer (50 mM Tris-HCl [pH 7.5], 150 mM NaCl, 1 mM EDTA, 1% NP-40, and 14.4 μM 2-mercaptoethanol), and incubated with a protease inhibitor cocktail (Roche, Basel, Switzerland) at 4 °C for 20 min. After centrifugation, the supernatant was mixed with sodium dodecyl sulphate (SDS) loading buffer and boiled for 5 min at 95 °C. After boiling, the samples were applied to a 4-15% polyacrylamide gradient/SDS gel (BIO-RAD, Hercules, CA). Proteins were electrophoretically separated and transferred to polyvinylidene fluoride (PVDF) membranes (Immobilon P, Merck Millipore, Burlington, MA, USA). Blocking treatment was performed using the PVDF Blocking Reagent for Can Get Signal (Toyobo) and the membranes were incubated with primary antibodies diluted with the Can Get Signal 1 reagent overnight. After incubation with secondary antibodies diluted with the Can Get Signal 2 reagent, the bands were visualized through chemiluminescence.

### Antibodies

The following antibodies were obtained commercially: anti-tRNase ZS (sc-101113, Santa Cruz, Dallas, TX, USA), anti-TRNT1 (NBP1-86589, Novus Biologicals, CO, USA), and anti-Vinculin (Proteintech, Rosemont, IL, USA).

### Viruses

TMEV (DA strain) and IAV (PR8 strain) were generous gifts from Dr. Yoshiro Ohara (Kanazawa Medical University, Ishikawa, Japan) and Dr. Toshiki Himeda (Kanazawa Medical University), and Dr. Mitsutoshi Yoneyama (Medical Mycology Research Center, Chiba University, Chiba, Japan), respectively. EMCV (VR-129B strain S) was purchased from the American Type Culture Collection (Manassas, VA, USA).

### Statistical analysis

Results were presented as mean ± standard deviation (SD). Differences between the control cells and the cells infected with EMCV were analyzed by Student t-test.

## Results

### tRNase ZS and TRNT1 mRNA levels are upregulated in cells infected with TMEV

We examined tRNase ZS and TRNT1 mRNA levels in HeLa cells after viral infection using qRT-PCR. HeLa cells were infected with TMEV at two different multiplicities of infection (MOI), normal MOI (MOI=1) and high MOI (MOI=10), and the mRNA levels were measured 10 and 16 h after the infection. The tRNase ZS and TRNT1 mRNAs were upregulated up to 13-fold and 3.6-fold, respectively, in 10 h, and neither mRNA was upregulated in 16 h (Fig. 1A and B). Upregulation of the tRNase ZL mRNA was not observed in either MOI (Fig. 1C). The infection of TMEV was confirmed by the upregulation of the interferon β (IFN-β) mRNA (Fig. 1D). The IFN-β mRNA was highly upregulated in a MOI-dependent manner (by 480- and 2800-fold) in 10 h, and its levels were drastically decreased in 16 h, resulting in reductions to 3 and 10%, respectively (Fig. 1D).

**Figure 1.**
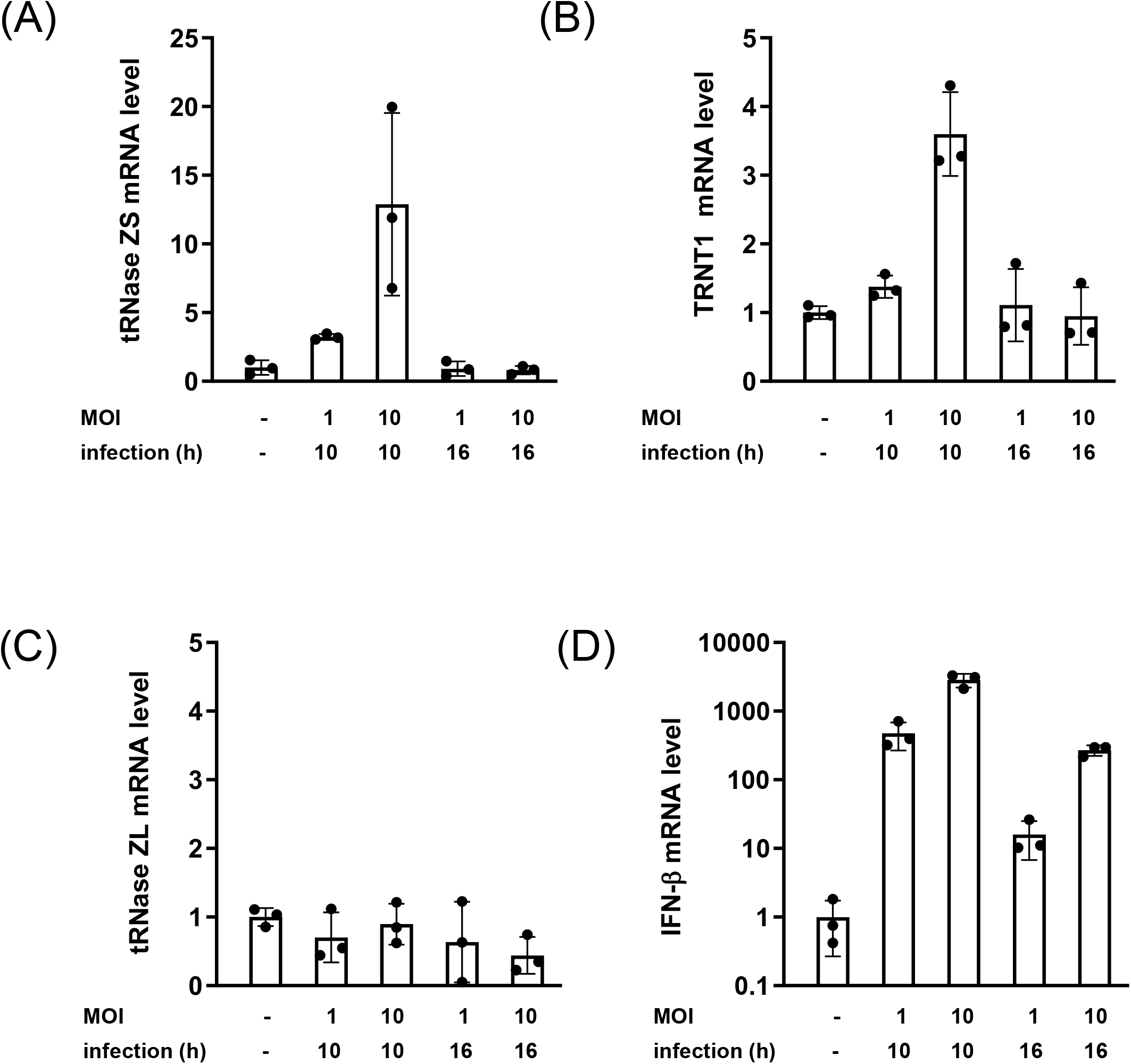
The mRNA levels of tRNase ZS and TRNT1 after infection with TMEV. The tRNase ZS (A), TRNT1 (B), tRNase ZL (C), and IFN-β (D) mRNAs were analyzed by qRT-PCR. HeLa cells were infected with TMEV at MOIs of 1 and 10. The mRNA levels are normalized against the GAPDH mRNA levels and expressed relative to those in the non-infected cells. Error bars indicate SD (n = 3).

### tRNase ZS and TRNT1 mRNA levels are upregulated in EMCV-infected cells with reduced efficiencies

Overall, both mRNAs were upregulated in a time- and a MOI-dependent manner in the EMCV-infected HeLa cells, albeit with reduced efficiency. The maximum upregulation of the tRNase ZS mRNA was by 2.7-fold in 16 h after the infection with MOI=10 (Fig. 2A), and the TRNT1 mRNA level reached to 2.2-fold in 10 h after the infection with MOI=10 (Fig. 2B). The tRNase ZL mRNA was not upregulated at all (Fig. 2C). The IFN-β mRNA was also highly upregulated like in the cells infected with TMEV, although its time- and MOI-dependency was not observed (Fig. 2D).

**Figure 2.**
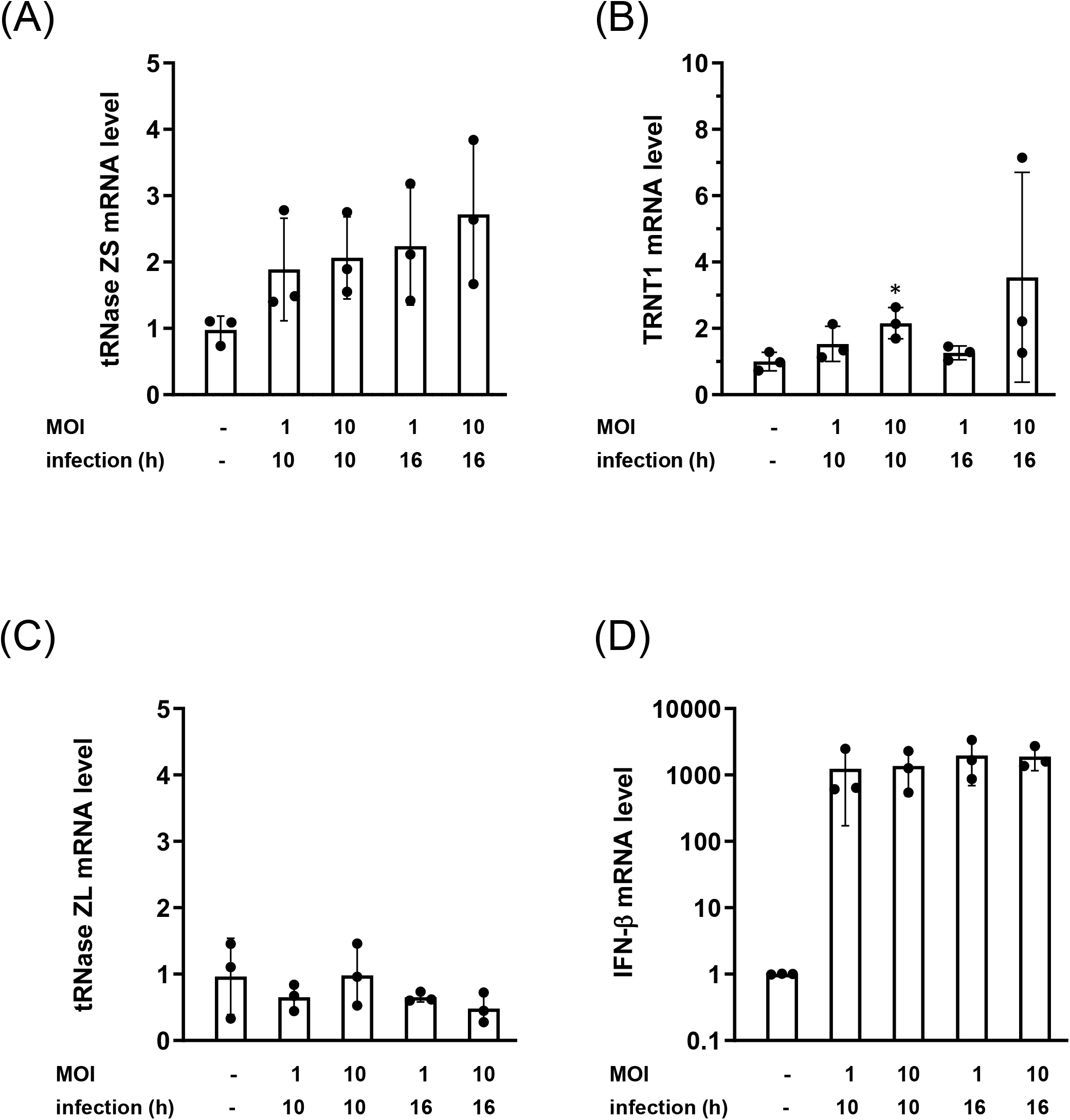
The mRNA levels of tRNase ZS and TRNT1 after infection with EMCV. The tRNase ZS (A), TRNT1 (B), tRNase ZL (C), and IFN-β (D) mRNAs were analyzed by qRT-PCR. HeLa cells were infected with EMCV at MOIs of 1 and 10. The mRNA levels are normalized against the GAPDH mRNA levels and expressed relative to those in the non-infected cells. Error bars indicate SD (n = 3). *P<0.05 vs. control.

### tRNase ZS and TRNT1 mRNA levels are not upregulated in cells infected with IAV

IAV neither upregulated tRNase ZS and TRNT1 mRNAs nor the tRNase ZL mRNA in the HeLa cells. (Fig. 3A–C). Since upregulation of the IFN-β mRNA was rarely observed following the IAV infection (data not shown), most likely due to interference by NS1 protein in IAV ^19^, its infection was confirmed through amplification of the mRNA encoding the IAV protein M. The protein M mRNA level in the IAV-infected HeLa cells was increased in a time- and a MOI-dependent manner (Fig. 3D).

**Figure 3.**
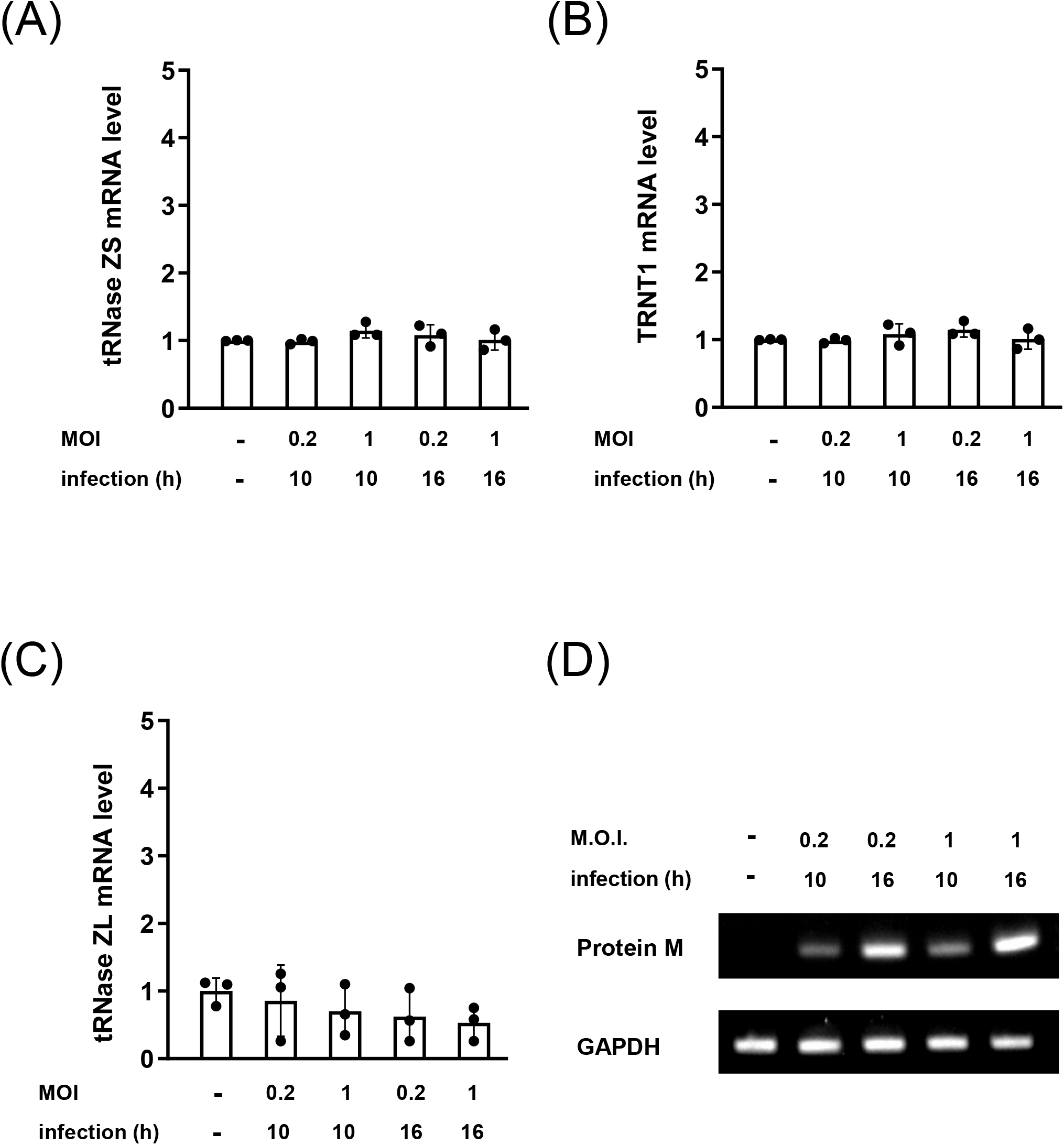
The mRNA levels of tRNase ZS and TRNT1 after infection with IAV. The tRNase ZS (A), TRNT1 (B), tRNase ZL (C), and IFN-β (D) mRNAs were analyzed by qRT-PCR. HeLa cells were infected with IAV at MOIs of 0.2 and 1. The mRNA levels are normalized against the GAPDH mRNA levels and expressed relative to those in the non-infected cells. Error bars indicate SD (n = 3). (D) RT-PCR products from the IAV protein M mRNA are shown. GAPDH cDNA was used as a loading control.

### tRNase ZS and TRNT1 proteins are upregulated in cells infected with TMEV

The tRNase ZS and TRNT1 protein levels after infection with TMEV were also examined. Although the expression pattern varied by experiments, on the whole, both protein levels after the MOI=1 infection increased up to 5 h and then decreased. And in the MOI=10 experiment, the tRNase ZS level reached a maximum in 3 h and the TRNT1 level continued increasing. A typical pattern is shown in Fig. 4.

**Figure 4.**
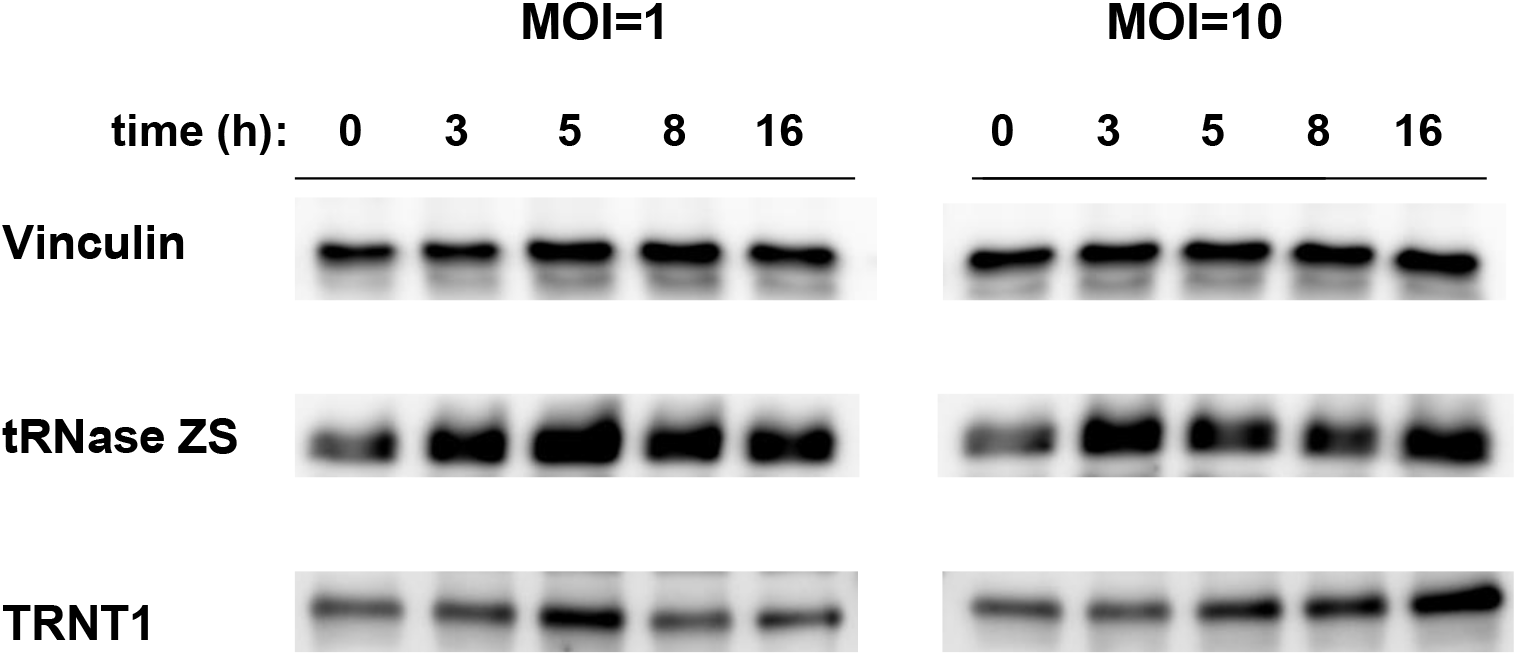
The protein levels of tRNase ZS and TRNT1 after infection with TMEV. HeLa cells were infected with TMEV at MOIs of 1 and 10. Whole-cell extracts were prepared 3, 5, 8, and 16 h after infection. Vinculin was used as a loading control.

## Discussion

Cellular responses to viral infection are mostly controlled by IFNs and more than 300 genes have been shown to be upregulated by them ^20,21^. Since the level of the IFN-β mRNA appeared to positively correlate with those of the tRNase ZS and TRNT1 mRNAs (Fig. 1–3), both genes may also be controlled by IFNs. Genes upregulated via the IFN-dependent pathway possess interferon <u>s</u>timulated responsible <u>e</u>lements (ISRE) and/or IFN-γ <u>a</u>ctivated <u>s</u>ite (GAS) elements in the promoter region ^22^. Further study from this aspect would elucidate a mechanism for tRNase ZS and TRNT1 gene upregulation under viral infection.

Although the tRNase ZS protein levels were also increased in the TMEV-infected cells, they were much lower than the levels expected from the mRNA levels (Fig. 1A and Fig. 4). This observation suggests that tRNase ZS may be regulated post-transcriptionally in the cells infected with TMEV and is consistent with our previous finding that *tRNase ZS* appears to be controlled post-transcriptionally in HEK293 cells^23^.

As far as we know, this is the first report showing that the mRNA levels of tRNase ZS and TRNT1, both of which are necessary for tRNA recycling, are upregulated by infection with TMEV or EMCV. We are currently trying to address the issues of whether tRNA recycling is indeed augmented under viral infection and, if so, what its physiological significance is.

## Conflict of interest

The authors declare no conflicts of interest.

## Acknowledgement

We thank Arisa Haino for her technical assistance. A research grant from the Niigata University of Pharmacy and Applied Life Sciences was received. The funding source was not involved in this study nor the publication of its results.

## Abbreviations

TMEV: Theiler’s mouse encephalitis virus
EMCV: encephalomyocarditis virus
IAV: influenza A virus
IFN-β: interferon beta
MOI: multiplicity of infection

## Notes

### Competing Interest Statement

The authors have declared no competing interest.

